# EEG-neurofeedback for promoting neuromodulation in the elderly: evidence from a double-blind study

**DOI:** 10.1101/2022.09.26.509227

**Authors:** Katia Andrade, Thomas Guieysse, Solofo Razafimahatratra, Nesma Houmani, André Klarsfeld, Gérard Dreyfus, Bruno Dubois, Takfarinas Medani, François Vialatte

## Abstract

**Background:** Electroencephalography (EEG) is a non-invasive method that records the brain signals with time resolution in the millisecond range, thereby allowing the monitoring of subjects’ mental states in real time. Using appropriate biomarkers extracted from these EEG signals and presenting them back in a neurofeedback loop can foster neural compensation mechanisms by teaching subjects to modulate their brain activity. Over the last decades, several neural biomarkers of aging have been described, with growing evidence suggesting that neuromodulation may have an important role in regulating brain activity in the elderly.

**Methods and objectives:** We used three neural biomarkers of aging, namely the Peak Alpha Frequency, the Gamma-band synchronization, and the Theta/Beta ratio, in the framework of an EEG-based brain-computer interface, with two main objectives: **1)** to test whether healthy elderly people with subjective memory complaints may learn to modulate their brain activity through EEG-neurofeedback training in a double-blind, placebo-controlled study; and **2)** whether that neuromodulation may have a positive impact on subjects’ cognition.

**Results:** A significant self-modulation of two neural biomarkers that usually decline with age was observed exclusively in the group of subjects who underwent the EEG-neurofeedback training, in clear contrast with the subjects who received the sham feedback. This neuromodulation did not have a direct effect on subjects’ cognitive abilities, as measured by neuropsychological tests pre-versus post-training, probably because all the subjects accurately performed these tests already at study entry (i.e., pre-training).

**Conclusion:** The results of this double-blind study are in line with one of the main criteria for successful neuromodulation, thus encouraging research on EEG-neurofeedback as a promising non-pharmacological intervention for promoting self-regulation of brain activity with a view to improve cognitive aging.

## 1 Introduction

Lifespan is increasing worldwide, which gives rise to new medical and socio-economic challenges. In this context, developing strategies for promoting successful aging has become a cornerstone of research in cognitive neuroscience.

Cognitive difficulties occur with aging even in the absence of brain disease. For instance, it is frequent that healthy elderly people present everyday memory complaints, such as forgetting where they put their glasses or their keys, which likely involves poor encoding and/or retrieval processes related to reduced attentional resources that mostly depend on the prefrontal cortex (Laborda-Sánchez et al 2021; Trambaiolli et al 2021; Hedden and Gabrieli 2004). This is important because it suggests that developing strategies for improving attention in the elderly may have a positive impact on healthy aging. Neurofeedbak is one such promising strategy (Ros et al. 2020; Niv 2013; Enriquez-Geppert, Huster, and Herrmann 2013). Specifically, neurofeedback training, through electroencephalography (EEG) passive brain-computer interfaces, has been found to improve attention in the elderly when focused on the Peak Alpha Frequency (PAF) – a neural biomarker of aging (Angelakis, Lubar, and Stathopoulou 2004; Angelakis et al. 2007; Wang and Hsieh 2013). Indeed, it is well documented that the PAF (i.e., the frequency of the spectral power density peak within the extended alpha band, 8 - 13 Hz) increases from early childhood until adolescence, remains stable during adulthood, and starts to decrease with age (Laborda-Sánchez et al 2021; Trambaiolli et al 2021; Scally et al. 2018). In patients suffering from either mild cognitive impairment (Garcés et al. 2013) or Alzheimer’s disease (AD) (Angelakis, Lubar, and Stathopoulou 2004), the PAF becomes pathologically lower than expected for healthy elderly people. Positive correlations between PAF and speed of information processing, PAF and memory performance, and PAF and inhibitory control have been reported (Bornkessel et al. 2004; Bazanova and Vernon 2014). Another neural biomarker of aging is the synchronization of neural activity in the gamma frequency band, which decreases with age, and even more in patients suffering from AD (König et al. 2005). This neural biomarker has been associated with several cognitive functions according to its role in neural systems for attention, memory, motivation and behavioral control (Bosman, Lansink, and Pennartz 2014). Additional neural biomarkers of aging include decreased EEG complexity, as well as an increased power of EEG low frequencies (i.e., delta and theta activities) together with a decreased power of high EEG frequencies (i.e., alpha, beta and gamma activities) (van der Hiele et al. 2007; Babiloni et al. 2016; Houmani et al. 2018).

The efficacy of EEG-neurofeedback can be measured based on cognitive outcomes (Laborda-Sánchez et al 2021; Trambaiolli et al 2021; Gruzelier 2014), but an obvious and more direct first outcome changes is at the level of brain activity itself (Orndorff-Plunkett et al. 2017). Indeed, it is hypothesized that EEG-neurofeedback can activate self-regulatory responses that can lead to the normalization of abnormal neural patterns. However, the lack of appropriate double-blind protocols has cast some doubts on the efficacy of neurofeedback interventions (Thibault, Lifshitz, and Raz 2016).

Here, we hypothesized that healthy elderly people could use EEG-neurofeedback training focused on neural biomarkers of aging in order to improve those biomarkers. To investigate, we transposed our biomarkers of interest – i) Peak Alpha Frequency; ii) Gamma-band synchronization; and iii) Theta/Beta ratio – to the framework of an EEG-neurofeedback system, and used a double-blind, placebo-controlled approach, with two main objectives: 1) to test whether elderly people may learn to self-modulate and improve their brain activity, based on specific changes in EEG neuronal dynamics pre-versus post-neurofeedback training; and 2) to test whether it may enhance their cognitive abilities as assessed by a battery of neuropsychological tests, mainly tapping into attentional, executive and episodic memory functions.

## 2 Methods

### 2.1 Participants

Thirty-seven subjects were recruited at the *Institut de la Mémoire et de la Maladie d’Alzheimer (IM2A)*, in the Salpêtrière hospital (Paris, France), where this double-blind study took place. The study was approved by the ethical committee and performed in accordance with the declaration of Helsinki. All subjects gave written informed consent prior to the experiments. Inclusion criteria were: cognitively healthy subjects older than 55 years presenting subjective memory complaints since at least six months, with a Mini-Mental State Examination [MMSE; (Burns, Brayne, and Folstein 1998)] score ≥ 27 and a Free and Cued Selective Reminding Test [FCSRT; (Grober and Buschke 1987)] total recall score ≥ 42.

Exclusion criteria were: concomitant neurological (neurodegenerative disorders, migraine, epilepsy, stroke, tumor) or psychiatric illnesses, as well as major visual disturbances, and the use of medications with a possible effect on cognitive performance, such as benzodiazepines, antidepressants, and antipsychotics. The eligibility criteria were verified by the medical investigator at the screening visit. This visit was also the pre-training visit (V0) for the subjects who met all the criteria. Further details are given in the next section.

### 2.2 Study design

This was a double-blind, placebo-controlled study, in which subjects were randomly assigned either to a real EEG-neurofeedback group (hereinafter called Group A) or to a sham feedback group (Group B), stratified by age, gender and education level. The allocation of the subjects to each of these two was performed by one of the collaborators not involved in the experiments, and remained hidde the investigators until the end of the data analysis.

In **Figure 1**, one can see a schematic representation of the study based on the timeline of the experimental protocol. The subjects were asked to participate in a research, neuromodulation protocol, over a three-month period (for each subject), during which they completed 22 visits: one pre-training visit (V0), 20 training visits (V1 to V20), and one post-training visit (V21). Neuropsychological and EEG data analyses performed in this study were based on the data collected during the pre-and post-training visits (V0 and V21) in order to investigate potential changes in the neural dynamics pre-versus post-training and their possible impact on subjects’ cognition.

**Figure 1.**
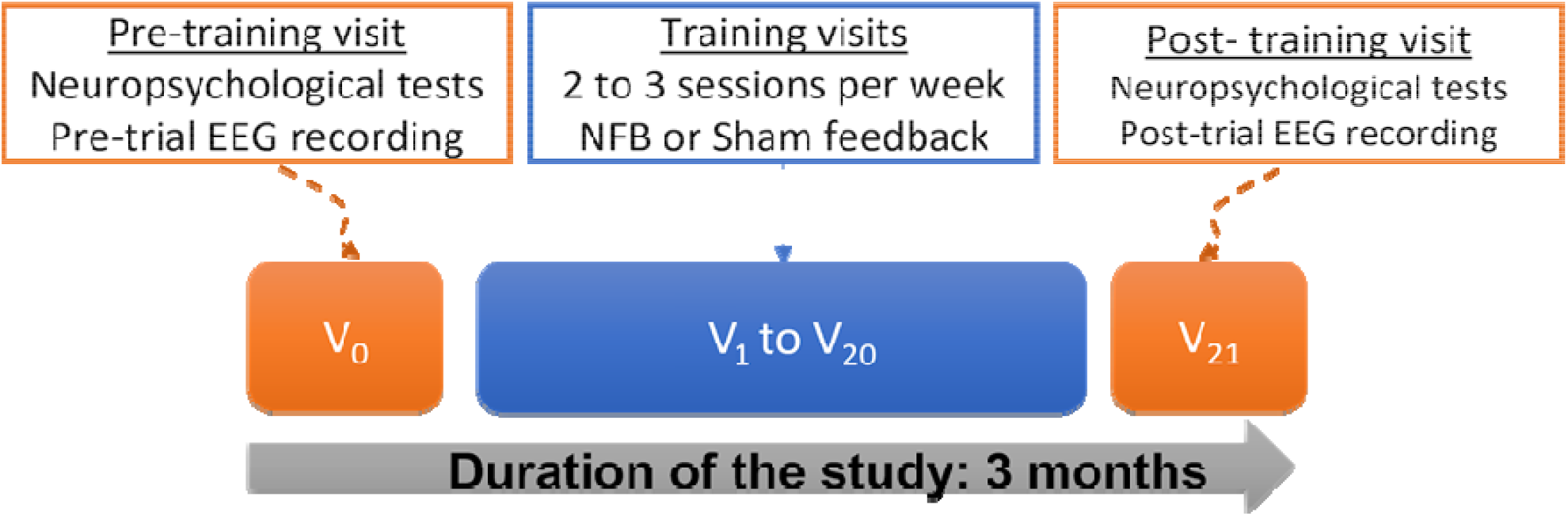
Schematic representation of the experimental protocol (NFB, neurofeedback)

For the neuromodulation, we built an EEG-neurofeedback brain-computer interface based on a small set of EEG electrodes in fronto-central and parieto-central sites (Fz, Cz, Pz, C3, C4) following anterior experiments that we had carried out on an EEG database of 22 subjects with Subjective Cognitive Impairment (SCI) and 28 mild AD patients. EEG data were recorded during resting state eyes-closed condition, using a Deltamed digital EEG acquisition system with 30 scalp electrodes, positioned over the whole head according to the 10-20 international system (Houmani et al. 2018). To discriminate between SCI and AD groups, we considered different combinations of electrodes among the 30 available, on which we computed an average Phase Synchrony measure at different frequency ranges. Then, such average value was considered as input of a Linear Discriminant Analysis classifier to evaluate the discrimination performance in terms of specificity (SCI correctly classified) and sensitivity (AD correctly classified). The best performance (specificity = 81.8% and sensitivity = 71.4%) were obtained using EEG electrodes of fronto-central and parieto-central sites.

#### 2.2.1 Neuropsychological assessment

The battery of neuropsychological tests administered to subjects included the MMSE (Burns, Brayne, and Folstein 1998), a measure of global cognition, and the FCSRT (Grober and Buschke 1987), a memory test that controls attention and cognitive processing to better investigate episodic memory performance, as well as tests focused on attention and executive functions (Dubois, Andrade, and Levy 2008), such as: the Trail-Making Test A and B (Reitan 1958); the Digit span forward and backward; and the Frontal Assessment Battery [FAB; (Dubois et al. 2000)]. Additional instruments included: the 15-item version of the McNair Frequency of Forgetting Questionnaire (McNair and Kahn 1983); the Geriatric Depression Scale [GDS; (Yesavage et al. 1982)]; and the State-Trait Anxiety Inventory [STAI A and B; (Spielberger 1983)] for assessing, respectively, memory complaints, mood, and anxiety disturbances.

#### 2.2.2 EEG data acquisition at V0 and V21

At V0, and upon completion of the neuropsychological tests, EEG data was acquired for 150 seconds (75 seconds eyes closed, and 75 seconds eyes open). A similar EEG acquisition was performed at V21, the post-training visit, for the subjects who completed the full training protocol. For both sessions (V0 and V21), the same EEG amplifier and same recording parameters were used: a 20-electrodes cap with a traditional 10-20 electrodes layout and a NIC® EEG amplifier system (Neuroelectrics, Barcelona, Spain) for recording the two blocks of EEG, 75 seconds with eyes open (looking at a fixation cross on the screen) and 75 seconds with eyes closed at rest. The left earlobe electrode was used as the EEG reference (Asper 1958). The impedance of all electrodes was kept below 10 kΩ. The sampling rate of EEG signals with the NIC system was 500Hz and no filter was applied during the acquisition phase. For the computation of our three biomarkers of interest, we used five electrodes on fronto-central and parieto-central sites: Cz, Pz, Fz, C3, and C4. Frequency intervals for the different biomarkers were considered as followed: Peak Alpha Frequency [8 – 13Hz]; Gamma-band synchronization [35 – 45Hz]; Theta/Beta ratio, respectively [4 – 8Hz] and [13 – 35Hz].

#### 2.2.3 Training visits between V1 and V20

The training started up to seven days after the screening, pre-training visit (V0): the feedback information was provided to each subject as a picture of a tree on her/his computer screen (see **Figure 2**). Subjects were only instructed to concentrate in order to make the tree grow. There were three different tree models: for each session, a different tree was assigned randomly to each biomarker. The training was performed in three blocks of 10 minutes per biomarker (30 minutes per session) during 20 visits over three months. At the beginning of each visit, one minute of the resting state EEG was recorded. These data were used to compute the threshold values of the subject’s biomarkers. During training, each biomarker was computed, in real-time, from the recorded signals. While the EEG-neurofeedback was estimated from the low-density EEG system based on the selected fronto-central and parieto-central electrodes, the sham feedback was estimated from the electromyography (EMG) sensors placed on the subjects’ trapezius muscles (two EMG sensors). If the actual computed value of the biomarker was higher than the threshold, the tree grew up; otherwise, the tree size remained constant. The design of the task was performed with the MATLAB Psychtoolbox (Brainard 1997) and the synchronization of the EEG and the stimuli presentation on the screen was performed with the SLS library (Kothe et al. 2014).

**Figure 2.**
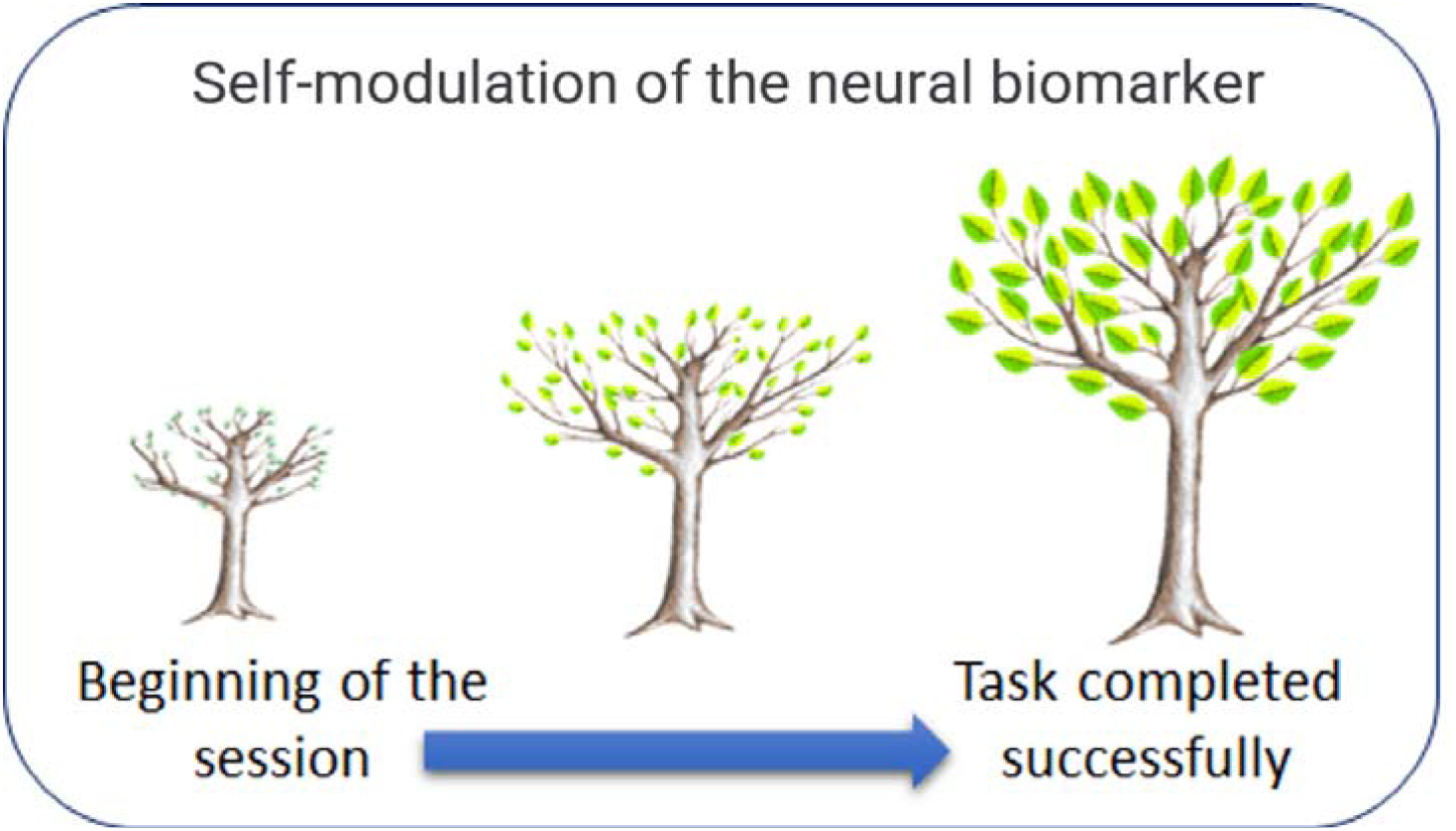
*Left:* Picture of two subjects during a session of the training phase. *Right:* The image of the feedback task is displayed on the screen. The subjects were asked to focus on modulating their mental states in order to grow the tree. Three different trees were used, with each neural biomarker being randomly assigned to one of the trees in each session. The sham biomarker was computed from the EMG sensors. The evolution of the tree on the screen is an indicator of subjects’ modulation of the neural biomarker.

### 2.3 Data analysis

#### 2.3.1 Demographic data

Thirty-seven subjects were initially included and randomly assigned into two groups, called Group A and Group B for preserving the double-blind approach. Six subjects desisted for personal, scheduling reasons at the beginning of the protocol, thus explaining the unbalanced number of subjects in the groups (17 subjects in Group A; 14 subjects in Group B).

Neuropsychological and EEG data analyses were performed on the data collected in pre- and post-training visits from the 31 subjects who completed the study. These subjects had a high education level and were all (except one) already retired. As summarized in **Table 1**, a significant difference in terms of gender ratio was observed between the two groups with a greater proportion of women in Group A (p = 0.05, Fisher’s test), while no significant differences were found between the groups for age and education level.

**Table 1.** Demographic characteristics of the subjects randomly distributed into two groups. p-values were computed using a Wilcoxon-Mann-Whitney test for the numerical variables (Age) and Fisher’s exact test for categorical variables (Gender, Education level). Significant differences are indicated by an asterisk. Categorizations of the education level into “high” and “low or intermediate” are as indicated in (Dubois et al. 2018). In short, more than 80% of our participants had an education level score ≥ 7, which corresponds to the label “high education” based on the (International Standard Classification of Education, 2011).

|  | Group A<br>(n=17) | Group B<br>(n=14) | p-value<br>(Intergroup A vs B) | All<br>(n=31) |
| --- | --- | --- | --- | --- |
| Gender<br>(M/F; % M, %F) | 4/13; 24% M, 76%F | 6/8; 43% M,<br>57%F | 0.05 * | 10/21; 33% M,<br>67%F |
| Age: mean ± SD<br>(range) | 71 ± 7.6<br>(60 - 85) | 73 ± 5.5<br>(64 - 82) | 0.44 | 72 ± 6.7<br>(60 - 85) |
| Education level<br>(% High) | 82% | 86% | 0.77 | 84% |

#### 2.3.2 EEG data pre-processing

The pre-processing of EEG data was performed on MATLAB version 2019, using the Brainstorm software (Tadel et al. 2011) in combination with custom scripts developed internally within the MATLAB environment (MathWorks). The sampling rate of the data during the acquisition was 500Hz, and no downsampling was applied in the pre-processing steps. The EEG signal was first band-pass-filtered between 0.5Hz and 45Hz to select frequencies corresponding to theta, alpha, beta, and gamma bands and remove the power line noise (50Hz). Data cleaning first included third-order spatial gradient noise cancellation, and Signal Space-Separation (SSP) to remove eye blink and saccade artifacts.

The filtered data, which had a total duration of 75 seconds per acquisition and per subject, were segmented into 2-second epochs. All the epochs with a kurtosis below 1 or above 5, or a peak-to-peak amplitude exceeding 100µV, were considered artifacts and rejected. The remaining epochs were visually checked, and the bad segments were identified and rejected. Due to the high number of artifacts on eyes-open data, the EEG analysis was performed only on the data collected from the eyes-closed condition (Hübner et al. 2018). The final number of epochs without artifacts per acquisition ranged from 32 to 37 for each subject. For the analysis, we decided to keep the same amount of data from each subject, thus retaining 32 epochs per subject.

Finally, values of the three biomarkers of interest were extracted from the pre-processed epochs. Then, two statistical approaches were conducted for data analysis, based on classical statistical tests and machine learning methods, as explained in the next two sections.

#### 2.3.3 Statistical tests on neuropsychological and EEG data

Since most neuropsychological and EEG data were not found to be normally distributed, non-parametric tests were performed in the analysis (for further details, see section 3). All statistical analyses were conducted using R version 3.6.1 [R Development Core Team, 2019] and plots were generated with the ggplot2 package (Wickham 2016). The level of statistical significance was set at p < 0.05 for all tests. To avoid type-I error, correction for multiple comparisons was completed by using the Bonferroni method (Armstrong 2014).

#### 2.3.4 Machine learning method for EEG data classification

We first define the term “feature”, which will be used throughout the rest of the paper. A feature is the value of one of the three biomarkers (PAF, Gamma-band synchronization, and Theta/Beta ratio), computed from an epoch of the EEG signal recorded in eyes-closed condition from a known electrode (PAF and Theta/Beta ratio), or from a known pair of electrodes (Gamma synchronization), during a known session.

In the present study, the features were computed from the measured EEG signals, segmented into 32 epochs as described in Section 2.3.2. Hence each subject contributed 64 examples (32 at session V0 and 32 at session V21) to the dataset available for classifier design. To train the classifiers, we used the SIGMABox (Medani et al. 2017), a homemade MATLAB toolbox for EEG signal processing and classification, to compute features from each of the three neural biomarkers using the EEG data.

Classifiers were designed for four different purposes: (i) discrimination of subjects of Group A from subjects of Group B, using the EEG epochs recorded during session V0; (ii) discrimination of subjects of Group A from subjects of Group B, using the EEG epochs recorded during session V21; (iii) discrimination of epochs recorded from subjects of Group A during session V0 from epochs recorded from the same subjects during session V21; (iv) discrimination of epochs recorded from subjects of Group B during session V0 from epochs recorded from the same subjects during session V21.

For all four purposes, the EEG signals were recorded from five electrodes (Cz, Pz, C4, Fz, C3). The PAF and Theta/Beta ratio biomarkers are computed from the signal captured by a single electrode; hence 5 features were computed from the PAF and 5 from the Theta/beta ratio. By contrast, Gamma-band synchronization requires two electrodes; as there exist 10 pairwise combinations of 5 electrodes, 10 features were computed from that biomarker. Thus, the classifiers had a maximum of 20 features. As all features may not be equally relevant for the classification, a subsequent feature selection procedure was applied to discard irrelevant features. The procedure has two steps: Orthogonal Forward Regression for ranking the features in order of decreasing relevance, followed by the random probe method to find the threshold rank for rejection of irrelevant features (Dreyfus et al. 2006).

For purposes (i) and (ii), the number of epochs available for the design of the classifier was 32 epochs per subject, hence 992 examples were available for classifier design. For purposes (iii) and (iv), the number of epochs available for the design was 64 epochs per subject, as the epochs to be classified were recorded during sessions V0 and V21, hence 1088 and 896 examples were available for classifiers (iii) and (iv), respectively. Table 2 summarizes the number of subjects and epochs.

**Table 2.** The number of subjects and of epochs in the training set and the test set of the classifiers.

|  |  | Training set<br>(70%) | Test set<br>(30%) | All data<br>(100%) |
| --- | --- | --- | --- | --- |
| Classifiers (i) and (ii) | Subjects | 22 | 9 | 31 |
|  | Epochs | 704 | 288 | 992 |
| Classifiers (iii) | Subjects | 12 | 5 | 17 |
|  | Epochs | 768 | 320 | 1088 |
| Classifiers (iv) | Subjects | 10 | 4 | 14 |
|  | Epochs | 640 | 256 | 896 |

The subjects of Group A and Group B (*n*=31) were randomly assigned to the training set (70%) and the test set (30%). As mentioned above, the classifiers were designed to perform epoch-wise classification from the features; the classification of subjects was performed according to the following rule: a subject was assigned to the class to which the majority of its epochs (at least 17) had been assigned by the epoch-wise classifier.

The classifiers were linear soft-margin Support Vector Machines (Burges 1998). The estimation of the regularization constant of the SVM was performed by leave-one-subject-out cross-validation. Due to the small size of the test set, the estimation of the performance of the classifier may vary with the partition of the examples into training set and test set. In order to decrease the variance of that estimation, cross-test was performed: instead of estimating the performance of the classifier on a single test set, 100 different partitions of the complete data set were performed, ascertaining that the number of inclusions *Ni* of each subject *i* in the test set was such that *5* ≤ *Ni* ≤ *8*. The whole design procedure was performed for each partition, thereby designing 100 classifiers. After completion of the iterations, the number of correct classifications for each subject, when he/she was present in the test set, was computed according to the above-mentioned subject classification rule. If the number of correct classifications of subject *i* was larger than the integer part of *Ni/2*, the subject was considered correctly classified. For instance, a subject with *Ni = 5* was considered correctly classified if he/she was correctly classified at least 3 times; a subject with *Ni = 8* was considered correctly classified if he/she was correctly classified at least 5 times.

## 3 Results

### 3.1 Statistical analysis of neuropsychological data

The results of the neuropsychological tests are detailed in the supplementary material (**Tables S1, S2, and S3)**. At study entry (V0), subjects’ performance across the entire battery of tests was within the normal range, and often close to the upper reference limit. For statistical analysis, nonparametric tests were used (_J = 0.05) because the criteria of normality were not verified for all the data. Regarding the TMT A test, we observed a significant improvement between V0 and V21 for the whole samples (subjects of Group A and Group B together) (*p* = 0.005), as well as for either group separately, being significant for Group A (*p* = 0.02) and a trend for Group B (*p* = 0.051). This suggests that subjects had a faster processing speed while performing this test at V21, regardless of the type of feedback training they received. Concerning the other neuropsychological tests, we did not observe any significant change between visits (V21 vs V0, intra-group analysis) nor between groups (Group A vs Group B, inter-group analysis) at V21, including also, in this case, the TMT-A. In fact, rather than suggesting a specific effect of feedback training, the improvement observed in the TMT-A test, at the post-training visit (V21), suggests a gain related to the previous exposure to this relatively simple attention test at V0, which is known as a practice effect. Regarding mood, anxiety, or memory complaints, an absence of significant differences was also observed in intra-group and inter-group analyses.

### 3.2 Statistical analysis of EEG results

#### 3.2.1 Inter-group analysis

##### Group A versus Group B at V0

To investigate whether the two groups (A and B) were different regarding their EEG activity at V0 (before the 20 training sessions), Mann-Whitney non-parametric tests were applied on the extracted 20 features (described in section 2.3.4). For each subject and each of the 20 features, a value was computed for each of the 32 epochs; then the mean of these 32 values was used as the biomarker associated to each subject. As shown in Table S4, irrespective of the biomarker or the EEG channel, there were no significant differences between groups at V0. This confirms that Group A and Group B were not significantly different regarding our three biomarkers of interest at the beginning of the study.

##### Group A versus Group B at V21

An inter-group analysis was also performed at V21 to determine whether the 20 training sessions had a significantly different effect on the EEG dynamics of the two groups (A vs B). Mann-Whitney non-parametric tests were applied on features (same procedure as presented in the previous section). As shown in **Table S5**, we observed a significant difference between groups regarding the Gamma-band synchronization for three pairs of electrodes: Pz/C4, Pz/Fz, and C4/Fz, with greater values on gamma-synchrony for Group A. A significant difference between groups was also found for this biomarker when considering the mean over all electrodes (*p* < 0.001). At V21, the groups were therefore significantly different concerning the Gamma-band synchronization, suggesting that subjects of Group A and Group B did not modulate this neural biomarker similarly post-training.

#### 3.2.2 Intra-group analysis

##### V0 versus V21 results, Group A and Group B separately

To investigate changes in the EEG data between pre- and post-training sessions (V0 vs V21), we compared, for each group, the value of our features at both V0 and V21. Thus, Mann-Whitney non-parametric tests were applied separately for each group (A and B), and each biomarker (i.e., PAF, Gamma-band synchronization, Theta/Beta ratio). As shown in **Table S6**, there were significant changes regarding the features computed for the Gamma-band synchronization for both Group A and Group B. However, for Group B, the difference was only observed in one pair of electrodes (Pz/Fz, *p* < 0.001), whereas, for Group A, a more global effect was corroborated by a significant increase of Gamma-band synchronization for the mean of all channels (*p* = 0.006). These results are consistent with the observation reported in the previous section, thus confirming that subjects of Group A and Group B did not modulate the Gamma-band synchronization similarly post-training.

### 3.3 Classification performance based on EEG data

As described in previous section, classifications were based on subjects’ epochs (epoch-wise classification). For inter-group comparison (Group A vs Group B, at V0 and V21 separately), a subject of the test set was considered as belonging to the group (A or B) to which more than 50% of her/his epochs were assigned. For intra-group comparison (V0 vs V21, Group A and Group B separately), the subject’s data of the test set was considered as belonging to the session (V0 or V21) to which more than 50% of her/his epochs were assigned.

#### 3.3.1 Inter-group classification at V0

In this section, the purpose of the classifier was to discriminate subjects of Group A from subjects of Group B based on their EEG data at V0. Using the three biomarkers, 20 features were extracted and the random probe method was applied to rank and select the most relevant ones. The first seven (out of 20) features were then used by the classifier to determine which epochs belonged to Group A or Group B.

As explained in section 2.3.4, variable selection was performed by (i) ranking the candidate variables in order of decreasing relevance by Orthogonal Forward Regression, and (ii) setting the variable rejection threshold by the random probe method. The random probe method consists in generating random variables (called probes) by shuffling the components of the vectors of the candidate variables, ranking the candidate variables and the probes in a single ranked list, and estimating the risk of selecting a candidate variable although it is irrelevant as the number of probe variables that rank better than the candidate variable. The higher that risk, the higher the number of selected variables, hence the higher the number of parameters of the model, hence the higher the risk of overfitting. A 20% risk of selecting a candidate variable although it is irrelevant turned out to be a reasonable choice, resulting in selecting the top 7 candidate variables in the ranked list and rejecting the others.

The **Table 3** shows the top 7 variables of a ranked list; with a 20% risk of accepting a variable although it is irrelevant (random probe variable p = 0.2), the first 7 variables were selected, and all others were rejected. The Gamma-band synchronization feature is predominant in the selected variables.

**Table 3.**
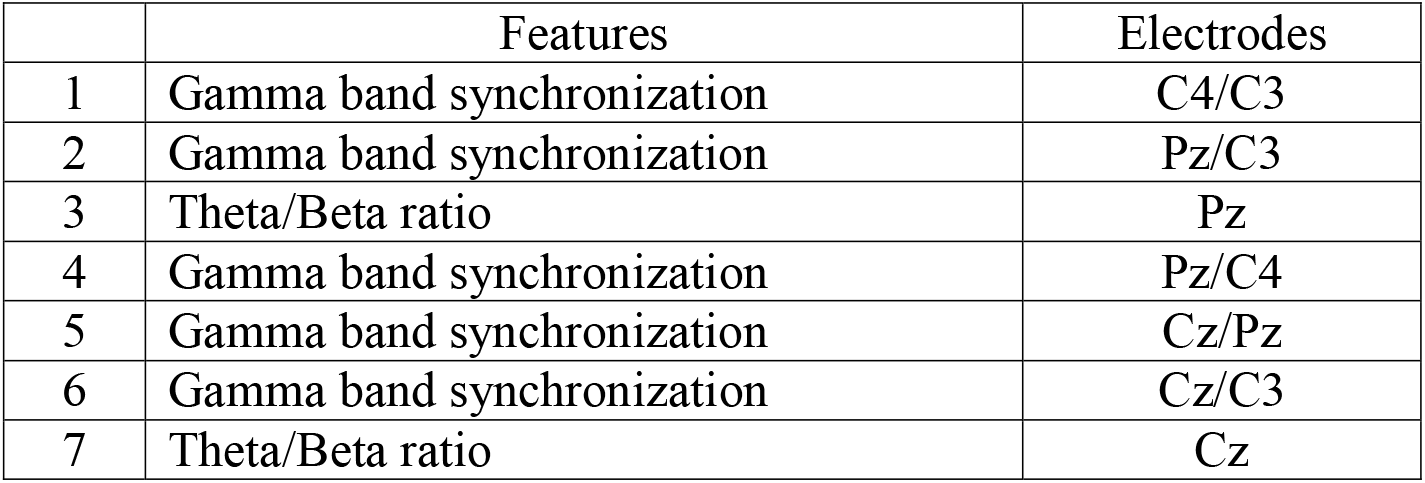
example of ranked list obtained by OFR; after application of the random probe method with a risk 0.2 of selecting a variable although it is actually irrelevant, the threshold of acceptance was set at 7: all variables ranking below the seventh variable of the ranked list were discarded.

**Table 4.** Confusion matrix resulting from the cross-test procedure on data recorded at V0

| Population (N = 31) | Group A | Group B |
| --- | --- | --- |
| Group A (N = 17) | 9 | 8 |
| Group B (N = 14) | 6 | 8 |

The following table shows the confusion matrix resulting from the cross-test procedure described previously. The accuracy of the classifier estimated by cross-test was 54.8%. It is good practice to compare the results of a classifier to the results that would have been obtained by a baseline classifier (called “zero-classifier”), which assigns all examples to the most populated class (Group A in our case), irrespective of the features; a classifier that does not provide a significantly better accuracy is unable to perform the classification. In the present case, the accuracy of our classifier is equal to that of the “zero-classifier”, which shows that the two groups cannot be discriminated on the basis of the three chosen biomarkers. This strongly suggests that the three EEG biomarkers computed from groups A and B were similar at V0, before training.

#### 3.3.2 Inter-group classification at V21

Here, the purpose of the classifier was to discriminate subjects of Group A from subjects of Group B based on their EEG data at V21. As explained in the previous section, the seven (out of 20) most relevant features selected by the random probe method were used to discriminate epochs recorded in subjects of Group A from epochs recorded in subjects of Group B. As presented in the confusion matrix resulting from cross-test (**Table 5)**, the classifier was able to classify data from Group A and Group B with a global accuracy of 71.0%. As the accuracy obtained is greater than 54.8% (the “zero-classifier”, see section 3.3.1), these results strongly suggest that the features computed from the three neural biomarkers were able to discriminate EEG data from groups A or B post-training.

**Table 5.**
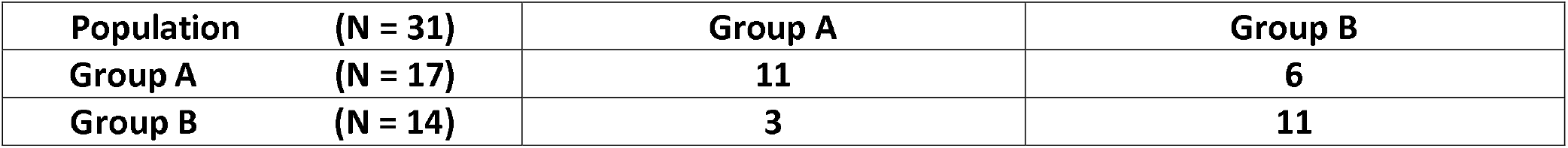
Confusion matrix of the classifier’s performance on the test data set at V21

| Population (N = 31) | Group A | Group B |
| --- | --- | --- |
| Group A (N = 17) | 11 | 6 |
| Group B (N = 14) | 3 | 11 |

Interestingly, at V21, among the 20 features extracted from the three biomarkers, the seven most relevant belonged in this case to the Gamma-band synchronization. Therefore, to validate the feature selection performed by the random probe method, three classifiers (i.e., one classifier per biomarker) were designed to perform the same inter-group analysis at V21. As presented in **Table 6**, the classifier whose features were extracted from the Gamma-band synchronization had an accuracy of 71%, while the accuracy of the classifiers trained with features extracted from either the Theta/Beta ratio or the Peak Alpha Frequency was 61.3% (estimated by cross-test). Interestingly, the accuracy of the classifier whose features were computed from the Gamma-band synchronization alone was equal to the accuracy of the classifier whose features were extracted from the three biomarkers simultaneously, with a satisfactory classification rate for both groups A and B (respectively, 12/17 and 10/14 well-classified subjects) at V21. This finding confirms the result of the feature selection by the random probe method, namely that the Gamma-band synchronization was particularly relevant to discriminate subjects of the two groups based on their respective epochs.

**Table 6.** True positive rate and accuracy, estimated by cross-test, of the classifiers discriminating subjects of Group A from subjects of Group B from each biomarker separately at V21. For more details, see supplementary material (Tables S7, S8, S9).

| Classification based on each biomarker | Gamma-band synchronization | Theta/Beta ratio | Peak Alpha Frequency |
| --- | --- | --- | --- |
| Group A ($n = 17$) | 12/17 | 11/17 | 11/17 |
| Group B ($n = 14$) | 10/14 | 8/14 | 8/14 |
| Accuracy | 71% | 61.3% | 61.3% |

#### 3.3.3 Intra-group classification (V0 versus V21)

Finally, we aimed to determine whether there was a significant difference between V0 and V21, within each group separately. In other words, we wondered whether subjects of Group A and Group B, separately, were able to self-modulate our biomarkers of interest post-training.

The classifiers using features extracted from the Gamma-band synchronization and the Peak Alpha Frequency were able to significantly discriminate between epochs recorded at V21 and epochs recorded at V0, only for subjects of Group A (accuracies of 67.6% and 61.8%, for the Gamma-band synchronization and the Peak Alpha Frequency, as shown in **Table 7**, versus accuracies of 35.7% and 32.1% regarding the same biomarkers in Group B, as shown in **Table 8**). By contrast, the classifiers using features extracted from the Theta/Beta ratio were unable to discriminate V0 from V21 in subjects of Group A (**Table 7**) and in subjects of Group B (**Table 8**).

**Table 7.** True positive rate and accuracy (estimated by cross-test) of the classifiers regarding the discrimination of the epochs recorded during V0 from those recorded during V21 from Group A, for each biomarker separately. For more details, see supplementary material (Tables S10, S11, S12).

| Group A | Gamma-band synchronization | Theta/Beta ratio | Peak Alpha Frequency |
| --- | --- | --- | --- |
| V0 ($n = 17$) | 11/17 | 6/17 | 10/17 |
| V21 ($n = 17$) | 12/17 | 8/17 | 11/17 |
| Accuracy | 67.6% | 41.2% | 61.8% |

**Table 8.** True positive rate and accuracy (estimated by cross-test) of the classifiers regarding the discrimination of the epochs recorded during V0 from those recorded during V21 from Group B, for each biomarker separately. For more details, see supplementary material (Tables S13, S14, S15).

| Group B | Gamma-band synchronization | Theta/Beta ratio | Peak Alpha Frequency |
| --- | --- | --- | --- |
| V0 (*n* = 14) | 6/14 | 7/14 | 5/14 |
| V21 (*n* = 14) | 4/14 | 1/14 | 4/14 |
| Accuracy | 35.7% | 28.6% | 32.1% |

As mentioned above, the fact that subjects who received a real, neural feedback belonged to Group A and subjects who received sham feedback belonged to Group B was disclosed after completion of data collection and analysis.

## 4 Discussion

One reason that healthy brain aging is emerging as a key issue for research in cognitive neurosciences is that the number of elderly people in populations around the world is increasing dramatically. The current study aimed to determine whether EEG-neurofeedback training (as opposed to sham feedback training) in healthy elderly people with subjective memory complaints could have a positive effect on both cognition and self-modulation of neural biomarkers of aging. There were no significant effects of the feedback training (either EEG-neurofeedback or sham feedback) on subjects’ cognitive abilities as measured by their performances in the neuropsychological tests. It is, however, possible that these subjects had a small room for improvement at V21 given that their performances in these tests were already within the normal range, and often close to the upper reference limit, at baseline (V0). The TMT A – a relatively simple timed test in which subjects are required to draw a line to connect consecutive numbers from 1 to 25 – was the only neuropsychological test in which the whole sample (n=31) or each group, separately, significantly improved their score at V21. However, rather than reflecting an improvement related to the feedback training, this finding suggests a processing-speed improvement probably due to practice. Importantly, the subjects’ education level was high, without significant differences between groups, which may have contributed to these cognitive results.

Concerning the EEG findings, the machine learning method not only confirmed, but further detailed the differences between biomarkers and between groups already verified with classical statistical tests, thus emphasizing the interest of that method in cognitive neurosciences. Among the 20 features extracted from our three neural biomarkers at V21, the SVM was trained with the seven most relevant: these features were able to accurately classify the epochs of 12 (out of 17) subjects in group A, and the epochs of 10 (out of 14) subjects in group B. The same accuracy was obtained by the classifier based on the features extracted from the Gamma-band synchronization alone. To further investigate the relevance of each biomarker, we performed intra-group analysis and compared, within-group A and within-group B, neural changes relative to each biomarker from V0 to V21. Our findings revealed that, at V21, the features extracted from the Gamma-band synchronization, as well as from the Peak Alpha Frequency, were able to correctly classify the epochs of subjects of group A (that is, epochs corresponding to 12/17 and 11/17 of subjects for the Gamma-band synchronization and the Peak Alpha Frequency, respectively), but not to discriminate the epochs of subjects of group B (that is, epochs corresponding to only 4/14 subjects, for either the Gamma-band synchronization or the Peak Alpha Frequency). Taken together, these results strongly suggest that subjects of group A, who underwent the EEG-neurofeedback training, were able to improve two neural biomarkers of aging (the Gamma-band synchronization and the Peak Alpha Frequency, but not the Theta/Beta ratio), while subjects of group B, who underwent the sham-feedback, were apparently unable to modulate any of these neural biomarkers. These findings are particularly interesting because they clearly distinguish Group A from Group B, based on EEG changes pre-versus post-training (V0 vs V21), which is in line with one of the main criteria for successful neurofeedback (Thompson 2003).

Overall, our findings, complementary statistical methods, and double-blind approach provide robust evidence that elderly people are able to self-modulate their brain activity, and specifically regulate two neural biomarkers of aging, through EEG-neurofeedback training.

## 5 Conclusion

The neuromodulation effect observed in this double-blind study constitutes a very encouraging finding in line with growing evidence suggesting that EEG-neurofeedback is a promising non-pharmacological intervention for improving brain activity in the elderly. This is particularly relevant with a view to maintaining a healthy brain during aging, because we know that, as the brain ages, the risk of developing neurodegenerative disorders like Alzheimer’s disease increases.

Future EEG-neurofeedback studies, specifically targeting the Gamma-band synchronization and the Peak Alpha Frequency, are required to replicate and validate our findings. Moreover, it would be of great interest to investigate whether these findings may extend to subjects with objective memory impairment, namely those suffering from AD at a prodromal stage (Dubois 2018), and further test whether, in this case, EEG-neurofeedback training could have an impact on patients’ cognition and quality of life by delaying AD progression.

## 7 Author Contributions

KA co-conceived the hypothesis, wrote and designed the protocol, and conducted the study, as the principal investigator; she further wrote the first draft and coordinated the final manuscript.

SR and TG participated in the acquisition of EEG data, performed behavioral and EEG analyses after the preprocessing step, performed the statistical analyses, discussed the results, and contributed to the text, tables and artwork of the draft and the final manuscript.

NH contributed to the design of the protocol and to the design of the EEG-based brain-computer interface, to the discussion of the results and to the writing of the final manuscript.

GD and AK contributed to the analysis and discussion of the results and to the writing of the final manuscript. GD further contributed to the preliminary statistical analysis.

BD contributed to the design of the protocol, to the discussion of the results and to the writing of the final manuscript.

TM co-designed and co-developed the task; participated in the acquisition of data and in EEG preprocessing; contributed to the discussion of the results and wrote the first draft and the final manuscript.

FV co-conceived the hypothesis and designed the study; contributed to the analysis and discussion of the results and to the writing of the final manuscript.

## 9 Acknowledgments

The authors are grateful to URGOTECH for their interest in our research, as well as to the research team of the IM2A, whose competencies on neuropsychological testing and subjects enrollment were particularly important for the success of this study. Also, we wish to thank all the subjects who participated in the current study for their strong commitment to this research.

## 10 Supplementary Material

Please refer to the following google doc: https://www.dropbox.com/s/f5tb1645np4pskx/Supplementary%20Material_for_submission.docx?dl=0

## 11 Data Availability Statement

The data generated and analysed in this study cannot be shared publicly. However, de-identified raw data could be shared under specific requests from the corresponding author [KA].

